# Life on the edge: microbial diversity, resistome, and virulome in soils from the Union Glacier cold desert

**DOI:** 10.1101/2024.05.13.594038

**Authors:** Patricio Arros, Daniel Palma, Matías Gálvez-Silva, Alexis Gaete, Hugo Gonzalez, Gabriela Carrasco, José Coche, Ian Perez, Jorge Gallardo-Cerda, Eduardo Castro-Nallar, Cristóbal Galbán, Macarena A. Varas, Marco Campos, Jacqueline Acuña, Milko Jorquera, Francisco P. Chávez, Verónica Cambiazo, Andrés E. Marcoleta

## Abstract

The high-latitude regions of Antarctica remain among the most remote, extreme, and least explored areas on Earth. Despite the highly restrictive conditions, microbial life has been found in these environments, although with limited information on their genetic properties and functional capabilities. Moreover, the accelerated melting of the Antarctic permafrost, the increasing exposure of soils, and the growing human transit pose the question of whether these environments could be a source of microbes or genes that could emerge and cause global health problems. In this line, although a high bacterial diversity and autochthonous multidrug-resistant bacteria have been found in soils of the Antarctic Peninsula, we still lack information regarding the resistome of areas closer to the South Pole. Moreover, no previous studies have evaluated the pathogenic potential of microbes inhabiting Antarctic soils. In this work, we combined metagenomic and culture-dependent approaches to investigate the microbial diversity, resistome, virulome, and mobile genetic elements (MGEs) in soils from Union Glacier, a high-latitude cold desert in West Antarctica. Despite the low organic matter content, diverse bacterial lineages were found, predominating Actinomycetota and Pseudomonadota, with limited archaeal and fungal taxa. We recovered more than 80 species-level representative genomes (SRGs) of predominant bacterial taxa and the archaeon *Nitrosocosmicus* sp. Diverse putative resistance and virulence genes were predicted among the SRGs, metagenomic reads, and contigs. Furthermore, we characterized bacterial isolates resistant to up to 24 clinical antibiotics, mainly *Pseudomonas*, *Arthrobacter*, *Plantibacter,* and *Flavobacterium*. Moreover, some isolates produced putative virulence factors, including siderophores, pyocyanins, and exoenzymes with hemolytic, lecithinase, protease, and DNAse activity. This evidence uncovers a largely unexplored resistome and virulome hosted by deep Antarctica’s soil microbial communities and the presence of bacteria with pathogenic potential, highlighting the relevance of One Health approaches for environmental surveillance in the white continent.

**HIGHLIGHTS:**

-Union Glacier soils host a microbial community dominated by bacteria, mainly from the phylum Actinomycetota.
-Archaea from the *Nitrosocosmicus* genus (family Nitrosphaeraceae) were ubiquitously detected.
-Although extreme and remote, these soils host multidrug-resistant and potentially pathogenic bacteria. Some were cultured and tested *in vitro*.
-Metagenomes and species-level representative genomes revealed diverse putative resistance and virulence genes.
-Part of the putative antimicrobial resistance genes and virulence factors could be associated with mobile elements in bacterial genomes.

## 1. Introduction

Antarctica hosts unparalleled natural landscapes where the geographical isolation and extreme climatic conditions, including low temperatures, limited organic nutrients, high salinity, low humidity, and a relatively high concentration of toxic metals, provide unique ecosystems for life to thrive, adapt, and evolve (Cowan and Tow, 2004; Vlček et al., 2017). Despite these harsh conditions, Antarctica hosts a remarkable microbial diversity, far exceeding that initially anticipated (Cowan, 2014; Lambrechts et al., 2019; Tindall, 2004). Moreover, this microbiota would have developed remarkable adaptations, including salient metabolic features and molecular mechanisms, to resist the harsh environment (Ramasamy et al., 2023). Thus, their study is valuable to expand our knowledge of on-the-edge life on Earth and possibly other planets (Cassaro et al., 2021; Correa and Abreu, 2020; Pacheco et al., 2019).

The Union Glacier, located in the Ellsworth Mountains of West Antarctica near the 80° S parallel, has been described as an extreme environment particularly challenging for life. In this nearly undisturbed region, the available climatic series (2008-2012) indicate a mean daily air temperature of −20.6 °C, with an absolute minimum of −42.7°C (August 2008) and an absolute maximum of 0.5°C (January 2010). Wind speed frequently overpassed ∼45 km/h, and the maximum recorded was ∼110 km/h (Rivera et al., 2014). Regardless of these highly freezing conditions, the null availability of liquid water and the high UV irradiance (Cordero et al., 2014), prokaryotic (Li et al., 2019) and fungal (Barahona et al., 2016) microbial life have been described in soil and sedimentary rocks from the Union Glacier region. However, there is still limited information about the diversity and structure of the resident microbial communities and a lack of representative genomes from this environment.

Previous studies in the Antarctic Peninsula soils showed a high prevalence of resistance to metals (Romaniuk et al., 2018) and different antibiotic classes (Marcoleta et al., 2022; Santos et al., 2022; Yuan et al., 2019) among autochthonous bacteria isolated from humanized or less-intervened areas, including isolates resistant to up to 10 antibiotics. Moreover, the analysis of soil metagenomes from different locations of the Antarctic Peninsula and surrounding islands showed a wide variety of antibiotic and metal resistance genes, part of them associated with mobile genetic elements (MGEs) potentially transferrable to other bacteria (Marcoleta et al., 2022; Santos et al., 2022; Yuan et al., 2019). This evidence agrees with other reports of antimicrobial resistance (AMR) in environments with minimal anthropogenic intervention (Pawlowski et al., 2016; Shen et al., 2019), indicating that resistance mechanisms would result from thousands of years of evolution (Allen et al., 2009; D’Costa et al., 2011). Additionally, the microbial communities from soils with minimum anthropogenic contamination should provide a potentially rich genetic resource to explore the spectrum of natural antibiotic mechanisms present before the introduction of antibiotics in medical or farming practice (Goethem et al., 2018). If this natural resistome could be a source of novel resistance genes among pathogens, it deserves further investigation.

Beyond the resistome, remote environments have also been recognized for their potential to host novel pathogens and virulence factor genes (VFGs) (He et al., 2022; Kim et al., 2022). Moreover, before Antarctica froze more than 10 million years ago, it had a temperate to sub-tropical climate with rainforest environments hosting several plant and animal species, including *Nothofagus* trees, dinosaurs, amphibians, and ancient mammals (Defler, 2019; Kloess et al., 2020; Mörs et al., 2020; O’Gorman et al., 2019; Vento et al., 2022). Thus, bacterial pathogens associated with these organisms likely existed in several areas of Antarctica, some potentially remaining under the ice for millions of years. Besides, horizontal transference of antibiotic resistance genes (ARGs) via MGEs (Partridge et al., 2018) could give rise to antibiotic-resistant pathogens.

Even though most of the previous studies regarding the Antarctic soil resistome have focused on different areas of the Antarctic Peninsula, scarce information is available regarding remote high-latitude areas, such as the Union Glacier, where environmental conditions are even more extreme. Thus, in this study, we combined culture-dependent and -independent approaches to compare the microbial diversity among different Union Glacier areas and reveal the bacterial community present in these habitats discoverable either as species-level representative genomes (SRGs) or as isolates in axenic culture. Then, we identified the most dominant ARG types and mechanisms of action, as well as the most prevalent VFs, in bacterial communities and the set of SRGs. Taking advantage of a collection of 51 isolates from Union Glacier soils, we developed phenotypic assays to reveal antibiotic resistance phenotypes and virulence factor production capabilities of a set of bacterial isolates, identifying multi-resistant strains and some exhibiting pathogenic potential. Our results highlight the importance of One Health-inspired studies to evaluate the possible role of remote environments in the emergence of new pathogens or virulence and resistance genes, with implications for global health.

## 2. Material and methods

### 2.1 Soil samples collection

The soil DNA samples were collected during the 58th Chilean Scientific Antarctic Expedition (ECA58) in November 2021. Three principal areas in the Union Glacier were sampled (Figure 1): Edson Hills (EH) (-79.8356, -83.6519), the border of Rossman Cove (RD) (-79.7929, -82.9096), and in a summit of Rossman Cove (RU) (-79.7921, -82.9854) southwest of Mount Rossman. Approximately 300 g of soil was collected for each sample after removing the surface layer using an aseptic metallic shovel previously cleaned with 1 % sodium hypochlorite, distilled water, and 70% ethanol. Samples were stored in sterile Whirl-Pak plastic bags and kept at -20 °C until processing.

**Figure 1.**
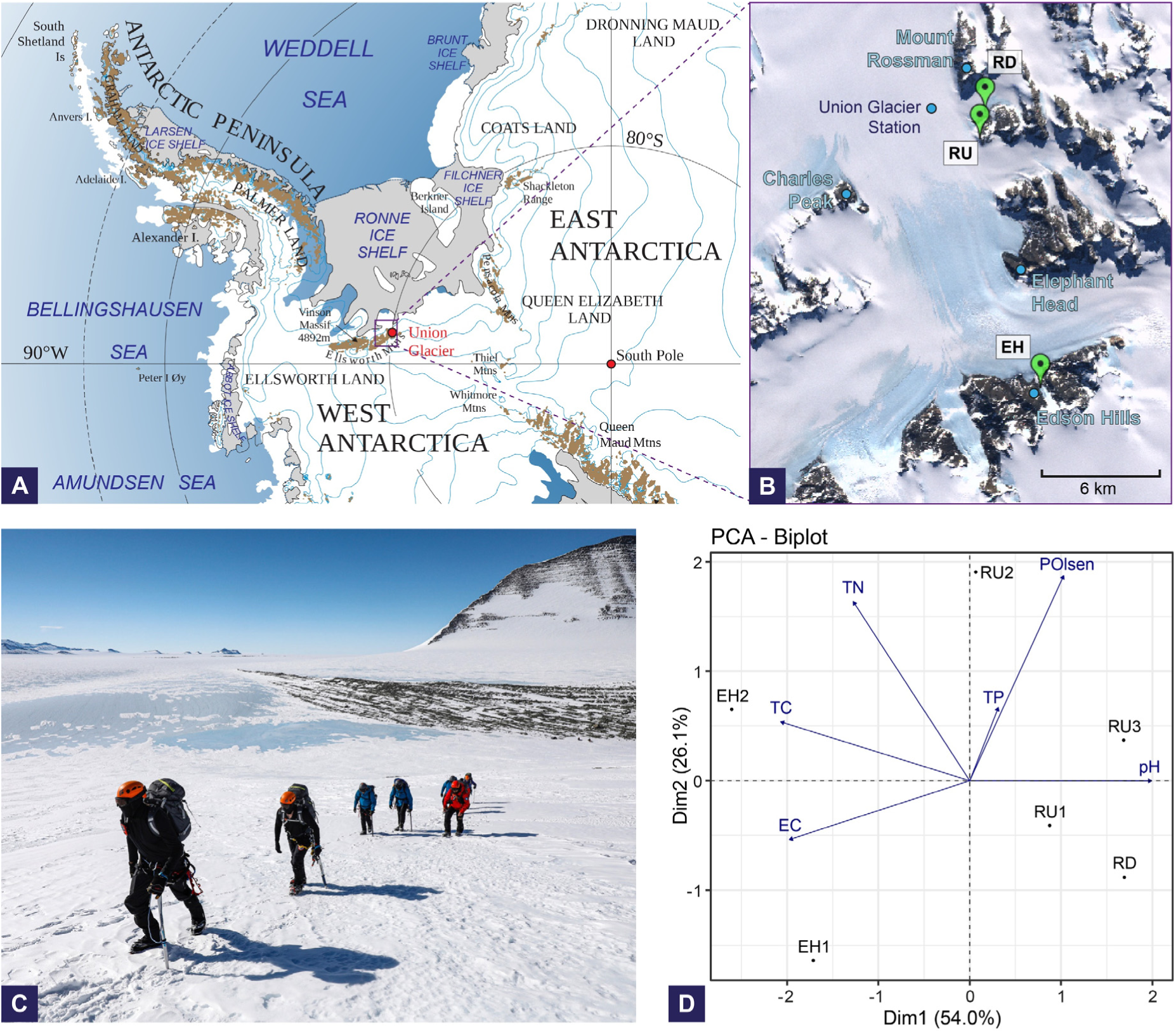
Geographical location of the sampled areas and distribution according to soil physicochemical properties. (A) Map of Antarctica showing the location of Union Glacier and the South Pole (red dots). Adapted from https://gisgeography.com/antarctica-map-satellite-image/. (B) Union Glacier area and the sampling points (green beacons; RD, RU, and EH). Satellite image obtained from SCAR with the respective GPS coordinates. (C) Photograph of the Union Glacier area landscape during the ECA58 Chilean Antarctic expedition. (D) Multivariate bidimensional projection of the sampled sites according to soil physicochemical properties.

### 2.2 Soil physicochemical parameters

The chemical properties of the soil samples (Supplementary Table 1) were determined as follows. Total carbon (TC) and nitrogen (TN) content were determined by loading freeze-dried and homogenized soil samples (150 µm pore size) on an automated elemental analyzer EA 3000 (Eurovector, Milano, IT), following the protocol described by Yang et al. (Yang et al., 2010). Organic matter (OM) content was estimated by wet digestion (Walkley and Black, 1934). The total phosphorous (TP) was determined by the ignition method, while the bioavailable phosphorous (POlsen) was determined following the Olsen method, based on 0.5 M sodium bicarbonate extraction. For both methods, the extracted P was analyzed using the molybdate-blue method (Murphy and Riley, 1962). Finally, pH and conductivity were determined in an aqueous solution (1:2.5 w/v) following the recommendations by Sadzawka R. (Sadzawka R., 1990).

### 2.3 Soil DNA isolation

Total DNA isolation was performed with the FastDNA Soil Kit (MP Biomedicals). Briefly, 500 mg of soil samples were introduced into Lysing Matrix E tubes (containing 1.4 mm ceramic spheres, 0.1 mm silica spheres, and one 4 mm glass bead). Subsequently, mechanical lysis was performed using a FastPrep homogenizer (MP Biomedicals) with three pulses at 6 m/s for 45 s, placing the samples on ice between each pulse. The following steps were made according to the manufacturer’s recommendations. The purified DNA was quantified by fluorometry using the Qubit dsDNA Broad Range (BR) kit (Invitrogen), and its integrity was corroborated by 1% agarose gel electrophoresis.

### 2.4 Metabarcoding sequencing

Three different molecular markers were amplified and sequenced using the Illumina NovaSeq system. For bacterial analysis, we selected the V3-V4 region of the 16S rRNA gene (primers 341F 5’-CCTAYGGGRBGCASCAG-3’ and 806R 5’-GGACTACNNGGGTATCTAAT-3’). For archaea, we targeted the V4-V5 region of the 16S rRNA gene (primers Arch519F 5’-CAGCCGCCGCGGTAA-3’ and Arch915R 5’-GTGCTCCCCCGCCAATTCCT-3’). Additionally, for fungal analysis, we targeted the region ITS2 (primers ITS3-2024F 5’-GCATCGATGAAGAACGCAGC-3’ and ITS4-2409R 5’-TCCTCCGCTTATTGATATGC-3’). Demultiplexed reads lacking primers, adapters, and barcodes were processed using Qiime 2 v2022.2 plugins (Bolyen et al., 2019). Amplicon Sequence Variants (ASVs) were generated using the paired reads DADA2 plugin (Callahan et al., 2016) and taxonomically assigned using the scikit-learn plugin with a naive Bayes classifier trained on the SILVA 138 database (Quast et al., 2013) for bacterial and archaeal ASVs, or the UNITE 9.0 database (2022-11-29) (Abarenkov et al., 2024) for fungal ASVs. The ASV alignments were carried out using MAFFT (Katoh and Standley, 2013), and FastTree was used to build the phylogenetic trees (Price et al., 2010). The phylogenetic tree visualizations were prepared using ETE 3 (Huerta-Cepas et al., 2016).

### 2.5 Diversity analysis

For diversity analyses, ASV abundance tables were first rarefied to 34500 counts per sample. Then, the R package vegan (Oksanen et al., 2022) was used to calculate the alpha diversity indices (Shannon index, Pielou’s evenness, and richness) for each sample, and the Bray-Curtis dissimilarity between samples.

### 2.6 Shotgun metagenomics

#### 2.6.1 Illumina and Nanopore shotgun sequencing

Shotgun Illumina sequencing was performed for RU and RD samples. RU1 and RU2 samples were pooled before sequencing. Due to insufficient metagenomic DNA, shotgun Illumina sequencing could not be completed for EH soils. Sequencing was carried out by Novogene Co (https://www.novogene.com/us-en/) on a NovaSeq platform (Illumina, San Diego, CA) with a 2 x 150 bp configuration, obtaining at least 12 Gb of data per metagenome. Illumina reads quality was checked with FastQC. Read filtering and trimming (to remove sequencing adapters and low-quality bases) were performed using fastp v0.23.2 (Chen et al., 2018).

Shotgun Nanopore sequencing could only be performed for the pooled RU samples, from which ≥1 µg of soil DNA was obtained. Library preparation was done using the Ligation Sequencing kit SQK-LSK110, which was then sequenced using a MinION Mk1c device and FLO-MIN106D flowcells. Basecalling was performed using Guppy v6.4.2. Nanopore read length and quality were checked using Nanoplot and filtered using Nanofilt, both from the NanoPack tools (De Coster et al., 2018).

#### 2.6.2 Sequence diversity and coverage assessment in shotgun metagenomic data

Possible human contamination of the metagenomic reads was removed by aligning the read sets against the GRCh38 human genome (used as a reference) with BWA-mem-2 v2.2.1 (Vasimuddin et al., 2019) and post-processing with SAMtools v1.16.1 (Danecek et al., 2021). Additionally, NonPareil v3.401 (Rodriguez-R et al., 2018) in kmer mode was used to estimate metagenomic coverage, sequencing effort, and alpha diversity (NonPareil diversity).

#### 2.6.3 Short-read-based virulence and resistance gene abundance calculation

ARGs and VFGs abundance was estimated using the metagenomic reads and the tool ARGs-OAP v3.2.3 with the SARGv3.2.1-S (Yin et al., 2023b) and VFDB (set B) (Liu et al., 2019) databases, respectively, following the consensus standardized pipeline proposed by Yin et al. 2023 (Yin et al., 2023a). ARGs-OAP was used with the default parameters, including the recommended cut-off values (E-value: 1e-7, identity: 80%, and hit length ratio: 75%) for positive hits when aligning the filtered reads against the proteins of the reference database using DIAMOND BLASTX. For metagenomic gene-of-interest (GOI) abundance calculation (ARGs and VFGs), the ARGs-OAP algorithm employs a function including the following parameters: 1) the number of GOI-like reads annotated to one specific GOI reference sequence, 2) the read length, 3) the nucleotide sequence length of the corresponding GOI reference sequence, 4) the number of mapped GOI reference sequences belonging to that GOI type or subtype (according to the respective database), and 5) the estimated total cell number in the metagenome. The cell number was estimated by mapping the reads against the essential single-copy marker gene database associated with ARGs-OAP (from the MicrobeCensus approach) (Nayfach and Pollard, 2015), with the default parameters. Heatmaps were constructed using ComplexHeatmap v2.16.0 package (Gu, 2022) in R.

#### 2.6.4 Metagenome assembly and SRGs recovery

Long ONT reads were removed from human contamination by aligning against the GRCh38 human genome reference with Minimap2 v2.24-r1122 (Li, 2021) and post-processed with SAMtools v1.16.1 (Danecek et al., 2021). The Illumina reads were assembled with metaSPAdes v3.15.5 (Nurk et al., 2017) with default parameters (Assembly A), while the ONT reads were assembled using Flye v2.9.1-b1780 (Kolmogorov et al., 2020) with default parameters (Assembly B). The hybrid metagenome assembly was performed using metaSPAdes v3.15.5 (Assembly C).

All the metagenomic assemblies obtained for each metagenome were *de novo* binned separately using MetaBAT2 v2.2.15 (Kang et al., 2019), AVAMB v4.0.0.DEV (Líndez et al., 2023), MetaCoAg v1.1.1 (Mallawaarachchi and Lin, 2022), SemiBin2 v1.5.1 in self-supervised mode (Pan et al., 2023), MetaDecoder v1.0.17 (Liu et al., 2022), and CONCOCT v1.1.0 (Alneberg et al., 2014). All binning processes were executed with default parameters and 139,808 as the initial seed. After all six *de novo* binning processes were obtained for each assembly (A, B, and C), ensemble binning was performed using DAS Tool v1.1.6 (Sieber et al., 2018) to recover initial MAGs. Then, initial MAGs were dereplicated into species-level representative genomes (SRGs) using dRep v3.4.3 (Olm et al., 2017) with a 98% ANI threshold and complete linkage.

#### 2.6.5 SRGs quality assessment, characterization, and phylogenetic analysis

SRGs were quality assessed with CheckM v1.2.2 (Parks et al., 2015), where Quality (Q) was defined as Q = Completeness - 5*Contamination. Taxonomic identification was performed using GTDB-tk 2.3.0 (Chaumeil et al., 2022) with database version 214. Only those SRGs with Q≥50% were kept for further analysis (medium to high-quality). The annotation, functional categorization of the encoded proteins, and visualization of the archaeal SRG (*Nitrosocosmicus* sp.) were done with the tool GenoVi (Cumsille et al., 2023).

A phylogenetic tree with the bacterial SRGs was constructed based on the multiple sequence alignment of 120 marker proteins obtained from GTDB-tk taxonomic identification using IQ-tree v2.1.4-beta (Minh et al., 2020) with default parameters (seed 139808). The highest likelihood amino acid substitution model was applied, Q.pfam+F+R8, as selected by the ModelFinderPlus algorithm (Kalyaanamoorthy et al., 2017). Tree visualization was done using iTOL v6.8.1 (Letunic and Bork, 2021).

The SRGs relative abundance in each metagenomic sample was determined by aligning the respective reads to the dereplicated SRGs using BWA-mem-2 v2.2.1 for short Illumina reads and Minimap2 v2.24-r1122 for long ONT reads. Read alignments were merged using SAMtools v1.16.1 for hybrid sequencing datasets. Next, read alignments were filtered stringently only considering alignments where at least 75% of the read was mapped to the reference sequence with 95% identity or more, following recommendations of Poff et al. (Poff et al., 2021) using CoverM v0.7.0 (https://zenodo.org/records/10531254). Filtered read alignments were then used to determine the SRGs’ relative abundance using CoverM v0.7.0.

#### 2.6.6 Identification of resistance and virulence genes in metagenomic assemblies and SRGs

All ORFs in metagenomic assemblies and SRGs were extracted using Prodigal v2.6.3 (Hyatt et al., 2010) in metagenomic mode and then aligned against SARGv3.2.1-L and VFDB (setB) databases using DIAMOND BLASTp v2.1.8 (Buchfink et al., 2021). Best hits with at least 50% identity, 50% bidirectional coverage, and between 0.8 to 1.2 length discrepancy (query length/subject length) were considered positive hits.

#### 2.6.7 Annotation of putative Mobile Genetic Elements carrying Genes of Interest in metagenome assemblies

Contigs containing positive hits against ARGs and VFGs were extracted from the assemblies and used as input for the prediction of complete integrons, clusters of attC sites lacking integron-integrases (CALIN), and integron integrases using IntegronFinder v2.0.2 (Néron et al., 2022). Integrated prophages were predicted using Phigaro v2.3.0 (Starikova et al., 2020). Complete and partial insertion sequences were predicted using ISEScan v1.7.2.3 (Xie and Tang, 2017). Predicted ISs located ≤5000 bp upstream or downstream from resistance or virulence genes were considered relevant hits. Filtered shotgun Illumina and ONT reads were assembled into putative plasmid sequences using metaplasmidSPAdes v3.15.5 (Antipov et al., 2019). Next, ARGs and VFGs were screened in assembled plasmids as described previously.

### 2.7 Isolation and taxonomic classification of soil bacteria

For bacterial isolation, the soil was mixed with sterile 1X PBS (1:1 ratio), and the suspension was mixed by vortexing for 40 min at maximum speed and then allowed to decant. 100 µL of the supernatant were plated on different agar media, namely, R2A (Difco, USA), Czapek (Difco), Nitrogen Free Medium (NFM) (HiMedia), and Luria Bertani (LB). The plates were incubated between 7 to 30 days at 4°C and 15°C. The obtained colonies were classified based on color, size, mucoviscosity, and microscopic features, including Gram staining and cell morphology examination. The isolates were first selected based on these criteria, discarding isolates exhibiting the same phenotypic features.

For taxonomic classification, the 16S rRNA gene for each isolate was amplified using the 27F (5’-AGAGTTTGATCCTGGCTCAG-3’) and 1492R (5’-GGTTACCTTGTTACGACTT-3’) primers and subjected to Sanger sequencing (Macrogen Inc.). The sequences were trimmed to eliminate low-quality nucleotide bases using the CLC software (version 12) and compared with the NCBI 16S ribosomal RNA sequences database using BLASTn (Altschul et al., 1990) (accessed on March 11^th^, 2024).

### 2.8 Antimicrobial sensitivity and virulence assays

The antimicrobial sensitivity of the selected isolates was tested using the disk diffusion method adapted to Antarctic bacteria, as previously described (Marcoleta et al., 2022). Thirty-five antibiotics representative of different classes were tested (Supplementary Table 2). Pre-cultures were set up in Mueller-Hilton (MH) broth and incubated for 48 h at 15 °C. Then, the cultures were adjusted at an optical density at 600 nm (OD_600 nm_) of 0.3 and used to inoculate MH agar plates using sterile swamps to complete a homogenous inoculum layer. The disks were applied with a 6-at-once disk dispenser (Oxoid) on the inoculated plates, which were then incubated at 15°C for 96 h. Antibiotic sensitivity was interpreted from the size of the growth inhibition halos, measuring the distance between the edge of the antibiotic disk and the bacterial growth area (<13 mm: resistant; >13 mm: sensitive).

Virulence factor production was evaluated by streaking the isolates on different agar media, namely, Egg Yolk, *Pseudomonas* CN (Winkler), DNAse toluidine blue (Winkler), and TSA with 5% sheep blood (Winkler). The inoculated plates were incubated at 15°C for 5-7 days, and the colonies’ growth and expected medium changes were evaluated. Also, we used the chrome azurol S agar (CAS), prepared as described previously (Schwyn and Neilands, 1987), but using R2A agar instead of supplemented M9. The R2A-CAS plates were inoculated with 10 µL bacterial suspension adjusted to OD^600^ = 0.3. Drops of suspensions prepared from six different Antarctic isolates and two control strains were inoculated per plate. We used the *K. pneumoniae* SGH10 Δ*wcaJ* strain, producing several siderophores (Tan et al., 2020), as a positive control. We used *E. coli* Δ*entB* from the Keio collection (Baba et al., 2006), which does not produce siderophores as a negative control. The presence of orange halos was evaluated and photographed after seven days of incubation at 15°C, protected from light.

## 3. Results

### 3.1 Soil physicochemical properties and microbial diversity

Soil samples were collected from three zones in Union Glacier: a summit of Rossman Cove (RU), a border of Rossman Cove (RD), and a summit of Edson Hills (EH), all part of the Elsworth Mountains (Figure 1A-C). The associations among sampled sites and their environmental variables (Supplementary Table 1) were explored in a multivariate space. As a result, the component that mainly explained the ordination of the sites along the transect (54%, strongly associated with electrical conductivity (EC), total cabron (TC), and pH) separated EH from the other soil samples (Figure 1D).

Amplicon sequencing of the 16S rRNA gene or the ITS region was used to evaluate the total bacterial, archaeal, and fungal diversity in the sampled soils. ASVs were classified into 17 bacterial, three archaeal, and two fungal phyla. Bacteria predominated in all samples, although a different community composition was observed when comparing samples from Edson Hills and Rossmann (RU and RD) (Figure 2A-B, Supplementary Table 3). In EH samples, the phylum Pseudomonadota (51.7% in EH1 and 93.4% in EH2) and the class Gammaproteobacteria were dominant, followed by the phylum Bacillota (19.7 in EH1 and 6.6% in EH1) and classes Bacilli and Clostridia (Figure 2A). The most abundant phyla in RD samples were Pseudomonadota (32%, mainly class Alphaproteobacteria), Acidobacteriota (23%, mainly class Blastocatellia), and Actinomycetota (37.2%, mainly classes Actinobacteria and Thermoleophilia). Similarly, RU samples were dominated by members of phyla Actinomycetota (41.6% in RU1, 36.3% in RU2, and 22.8% in RU3), followed by Acidobacteriota (45.4% in RU1, 35.9% in RU2 and 23.2% in RU3), with presence of the classes Actinobacteria, Thermoleophilia and Blastocatellia (Figure 2A-B, Supplementary Table 3).

**Figure 2.**
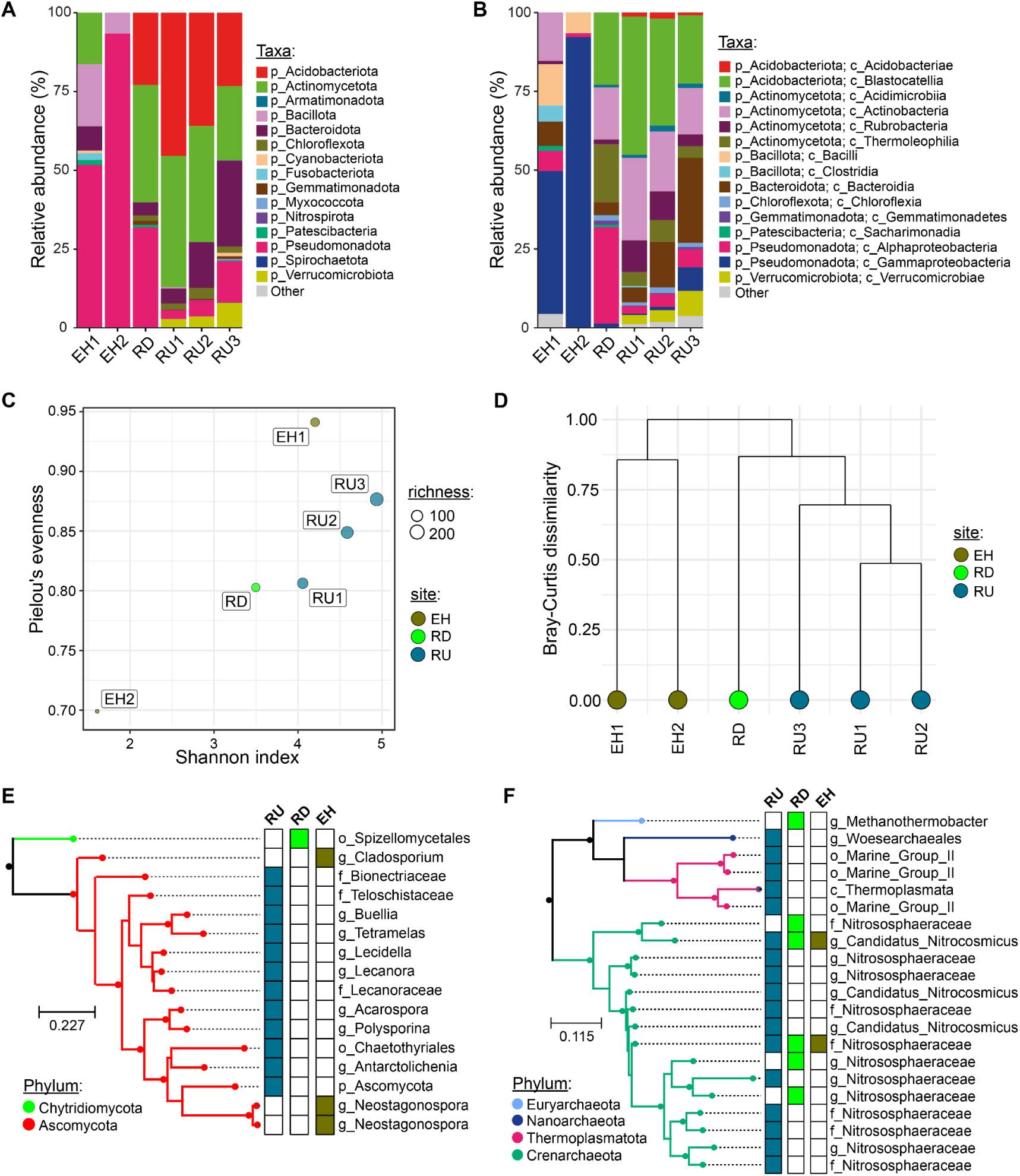
Bacterial, fungal, and archaeal diversity in the Union Glacier sampled soils. (A) Relative abundances of bacterial taxa at the phylum level. (B) Relative abundances of bacterial taxa at the class level. (C) Alpha diversity metrics of bacterial ASVs. (D) Complete linkage hierarchical clustering based on the Bray-Curtis dissimilarity matrix of bacterial ASVs. (E) Phylogenetic tree of fungal ASVs. (F) Phylogenetic tree of archaeal ASVs.

Bacteria of two genera, *Pseudomonas* and *Acinetobacter*, dominated the two samples from EH, whereas the most abundant genera in RD samples were *Blastocatella*, *Nocardioides,* and *Sphingomonas*. A markedly higher abundance of *Blastocatella* was observed in the three samples from RU, followed by *Nocardioides* and *Rubrobacter*. A detailed view of the phylogenetic relationships among the ASVs and their abundances in the different sampled soils is depicted in Supplementary Figure 1, while the complete list of microbial ASV counts per sample is provided in Supplementary Table 3.

Apparent differences in alpha diversity (Shannon index and evenness) were detected between the soils of the Union Glacier; in particular, the EH2 microbial community exhibited the lowest diversity, mainly due to a limited evenness, in agreement with the high predominance of *Pseudomonas* in this sample (Figure 2C). Additionally, beta-diversity analysis by hierarchical clustering based on the Bray-Curtis dissimilarity indicated that RD and RU samples had a similar bacterial community structure while EH samples clustered apart (Figure 2D), which agrees with the relative abundances of bacterial taxa observed.

Fungal ASVs exhibited very limited richness in these soils. In RU and EH samples, we detected 15 ASVs affiliated with the phylum Ascomycota and the classes Lecanoromycetes and Dothideomycetes. In RD, only one ASV from the phylum Chytridiomycota and the order Spizellomycetales was detected (Figure 2E, Supplementary Table 3). The members of the Ascomycota phylum belonged to 12 genera, including *Cladosporium* and *Neostagonospora* from EH1 and EH2, respectively, and *Lecidella* detected in RU1 and RU3 samples. On the other hand, the RU2 sample showed the highest number of ASVs, belonging to genera *Acarospora*, *Buellia*, *Antarctolichenia*, *Polysporina*, *Tetramelas,* and *Lecanora*.

Archaea found within the Union Glacier soils were limited to Crenarchaeota, Thermoplasmatota, Euryarchaeota, and Nanoarchaeota phyla (Figure 2E, Supplementary Table 3). The most significant number of archaeal ASVs (N=17) was detected in the RU soil, and most of them were affiliated with the family Nitrosphaeraceae (Crenarchaeota), which was the only archaeal taxon detected in the three sampled soils (RU, RD, and EH). The rest of the archaeal ASVs belonged to the genus Methanothermobacter (Euryarchaeota), order Woesearchaeales (Nanoarchaeota), or were affiliated with the Marine Group 1.1b (Thermoplasmatota).

### 3.2 Union Glacier microbial communities as revealed by shotgun metagenomics and species-level representative genomes

Next, we used shotgun metagenomics to gain deeper insights into the Union Glacier microbiota’s diversity and genomic features. RU and RD samples, but not EH samples, yielded enough DNA for Illumina shotgun sequencing, while we could also perform Nanopore sequencing for RU. A total of ̴19 and ̴ 16 Gbp of Illumina reads were obtained for RU and RD, respectively, plus ̴ 16 Gbp of Nanopore reads for the former (Supplementary Table 4). Metagenome coverage and sequence diversity analysis using Nonpareil indicated 79% and 88% coverage, with Nonpareil diversity values of 21.1 and 19.7, in the range of low-complexity soils (Rodriguez-R et al., 2018). This relatively low sequence diversity agrees with the metabarcoding analyses, pointing out that only a limited number of taxa can thrive in the extreme environmental conditions occurring in these soils.

Metagenome assembly for RD (Illumina-only) yielded 404.5 Gbp with an N50 of 2.97 kbp (Supplementary Table 4). We used both read sets (Illumina and Nanopore) for RU to build three assemblies (Illumina-only, ONT-only, and hybrid). ONT and hybrid assemblies had three times more assembled bases than Illumina-only (∼1.1-1.2 Gbp versus ∼480 Mbp), with notably higher contiguity, showing an N50 value of 56.8 and 5.59 kbp, respectively. Thus, introducing the long Nanopore reads notably impacted the assembly quality. After comparing the three RU assemblies using Mash distance and ANI calculations, we noticed they had slightly different sequence compositions and thus likely captured complementary genetic information. Therefore, we proceeded with the binning and genome reconstruction, starting in parallel with all the assemblies and then pooling and dereplicating the bins into species-level representative genomes (SRGs), using a 98% ANI threshold.

A total of 83 medium- to high-quality SRGs were obtained (33 from RD and 50 from RU), with completeness ranging from ∼51.5 to 99.7% and contamination from 0 to 9.8% (Supplementary Table 5). To evaluate the taxonomic groups captured in our SRG set and their phylogenetic relationships, we performed genome-based taxonomic assignments using GTDB-Tk. Only one (nearly complete) archaeal SRG was recovered (RUP31), corresponding to *Nitrosocosmicus sp*. (Supplementary Figure 2), in agreement with the metabarcoding data indicating this genus belonging to the dominant phylum Crenarchaeota was present in the three Union Glacier sampled areas. The other SRGs encompassed eight bacterial phyla, mainly Actinomycetota (39), Bacteroidota (15), Chloroflexota (9), and Acidobacterota (8). Conversely, despite being present in the soil samples according to the metabarcoding analysis, no SRGs belonging to Bacillota, Cyanobacteriota, Verrucomicrobiota, and Deinococcota could be recovered, probably due to their lower abundance.

At least eight SRGs with potential taxonomic novelty were identified, outstanding five Actinomycetota, including two putative novel genera from the Nocardioidaceae family (RUP16 and RUP39), two novel species from the JACCRW01 uncultured genus (family Gaiellaceae) (RDP17 and RUP48), and a novel family from the order Acidimicrobiales (RDP23). Also, we obtained a representative genome of a putative novel genus from the Gemmatimonadaceae family (RUP05, phylum Gemmatimonadota), a novel species from the genus *Singulisphaera* (Planctomycetota, RUP30), and a novel species from *Luteimonas* (Pseudomonadota, RUP04). Overall, this SRG collection corresponds to valuable baseline knowledge regarding the genomic features of the microbiota living in this hyper-extreme environment and would serve future studies focused on their metabolic capabilities and biotechnological potential.

### 3.3. Virulome and resistome from Union Glacier soils

Next, we investigated the presence of putative antibiotic resistance and virulence genes in the microbiota living in Union Glacier soils. First, we identified antibiotic resistance and virulence genes using the Illumina reads, calculating and comparing their relative abundance in the two metagenomes (RU and RD). To this end, we used the ARGs-OAP tool (and the SARG database), specially designed and optimized for metagenomic ARG prediction and normalized gene abundance calculations (Yin et al., 2023b).

More than 45 antibiotic resistance gene families, including several efflux pumps and antibiotic inactivation enzymes, were identified, potentially conferring resistance to 16 drug classes (Figure 3A). The gene abundances ranged from zero to 0.03 copies per cell. Novoviocin resistance gene *novA* was the most abundant ARG in RD and had a high abundance in RU. For most of the other resistance gene families, we observed a greater abundance in RU compared to RD, including orthologs of the Mex efflux pumps, conferring resistance to multiple drugs, nine predicted beta-lactamase families, three different aminoglycoside acetyltransferases, and a set of putative rifampin ADP-ribosyltransferases (Arr proteins). Other resistance genes found in high abundance were *bacA* and *bcrA*, linked to bacitracin resistance, and genes involved in lipopolysaccharide modification (*arnA*, *arnC*, *pmrE*) related to polymyxin resistance.

**Figure 3.**
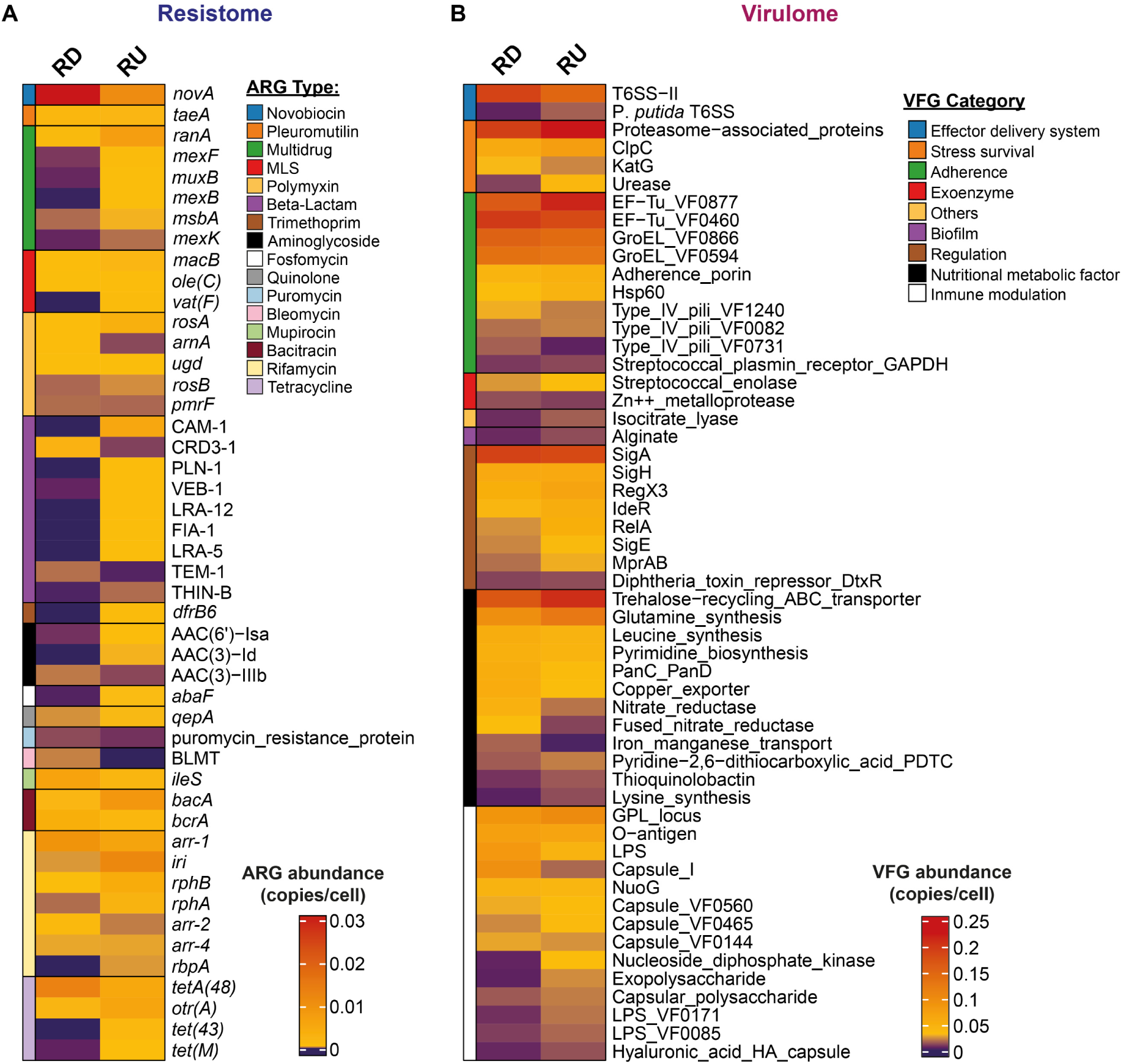
Metagenome-wide resistome and virulome from Union Glacier soils. The heatmaps show the relative abundance of genes encoding putative antibiotic resistance (A) and virulence determinants (B) in gene copies/cell, based on the analysis of unassembled Illumina reads. The ARGs and VFGs were classified according to the SARG and VFDB databases, respectively. The genes shown correspond to the virulence: 90% sum of the abundances in RU and RD; resistance: 95%.

We also used the Illumina reads to search for genes encoding putative virulence factors, using ARGs-OAP with the Virulence Factors Database VFDB. A relatively higher abundance of putative virulence factors was found in RU, with values up to 0.25 copies/cell. Among them, we identified genes linked to effector delivery (Type-6 secretion systems), stress survival (chaperones, catalases, ureases), adherence (Type-IV pili, GroEL), biofilm formation (alginate synthesis), exoenzymes (Zn++ metalloprotease, enolase), immune modulation (capsule synthesis, LPS), and nutritional metabolic factors (thioquinolobactin siderophore, amino acid and lipid metabolism, metal tolerance and metabolism) among other functions (Figure 3B). A significant proportion of the factors found are similar to genes commonly found in *Mycobacterium*, which agrees with the high proportion of Actinomycetota found in these soils. Also, several factors showed similarity to *Pseudomonas* genes.

### 3.4. Genomic context of the Union Glacier resistance and virulence genes

We next searched for putative virulence and resistance determinants among the recovered SRGs to gain further details about these genes, their genomic contexts, and the microbial taxa hosting them. We first examined the distribution of these genes across different phylogroups, as shown by a maximum-likelihood tree constructed based on the alignment of 120 ubiquitous bacterial marker genes (Chaumeil et al., 2022), matched with the antibiotic resistance and virulence gene prediction (Figure 4).

**Figure 4.**
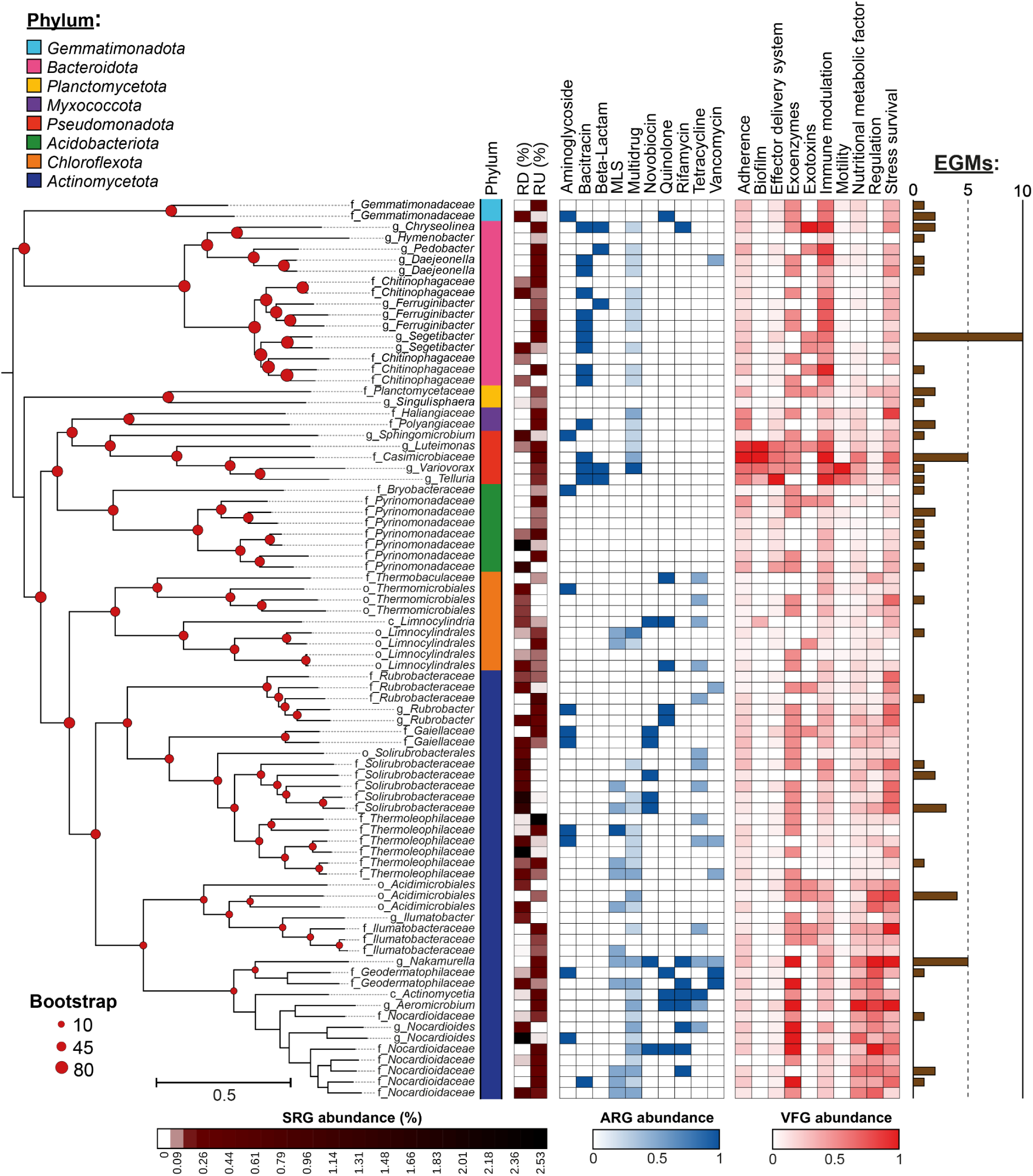
Species-level representative genomes recovered from Union Glacier soil metagenomes and their predicted virulence and antibiotic resistance genes. Maximum likelihood tree showing the phylogenetic relationships among 82 bacterial SRGs, inferred from the multiple alignments of 120 marker proteins as defined by the GTDB taxonomy. The bootstrap values were calculated from 1000 iterations. The taxonomic classification at the lowest rank according to GTDB is provided for each SRG. The abundance of each ARG and VFG category in each SRG was calculated as the number of hits per category divided by the maximum number of hits for that category across all the SRGs. The SRG abundance was estimated using the CoverM tool based on mappings of the short and long reads to each SRG. The mobile elements value corresponds to the sum of the insertion sequences, prophages, and integrons.

We calculated the abundance of a resistance or virulence gene class (e.g., “beta-lactam” or “adherence”) in a given SRG as the sum of hits from this class in the SRG divided by the highest number of hits of the same class observed in any SRG. From this, gene class abundance in a given taxon corresponds to the sum of the gene class abundance of all SRGs of the same taxon divided by the number of SRGs belonging to that respective taxon. Then, the highest total ARG abundance was observed in Pseudomonadota (1.60; mainly Burkholderiales, Sphingomonadales), followed by Actinomycetota (1.35; mainly *Mycobacteriales* and the uncultured JACCYY01 and Gaiellales), Bacteroidota (1.15; Cytophagales and Sphingobacteriales), Chloroflexota (1.00; QHBO01), and Gemmatimonadota (1.00; Gemmatimonadales). However, it varied depending on the resistance gene family (Figure 4 and Supplementary Table 6). In this line, novobiocin resistance was highly abundant in Actinomycetota, in agreement with *novA* being the most abundant ARG in RD and this phylum the most abundant in this soil sample. Actinomycetota SRGs also had the highest abundance of predicted ARGs to aminoglycosides (several acetyltransferases), MLS, rifamycin, and vancomycin. Meanwhile, the highest abundance of beta-lactam (including putative beta-lactamases LRA, CAM, and LUS) and multidrug resistance genes was observed in Pseudomonadota, followed by Bacteroidota SRGs.

For bacitracin, where the *bacA* and *bcrA* genes were also abundant in the two soil samples, the highest abundance is observed in Bacteroidota, followed by Pseudomonadota. For quinolones and tetracycline, the highest abundance was observed in Chloroflexota, followed by Actinomycetota.

Regarding virulence genes, as for ARGs, we confirmed the presence of a highly similar set of genes to those detected in the reads-based analysis (Figure 4, Supplementary Table 7). The highest total VFG abundance was observed in Pseudomonadota (4.36; Xanthomonadales, Burkholderiales), followed by Actinomycetota (2.13; Mycobacteriales, Propionibacteriales), Gemmatimonadota (2.04; Gemmatimonadaceae), and Myxococcota (2.01; mainly Haliangiales). Pseudomonadota showed the highest abundance of genes for adherence (mainly Type-IV pili and GroEL), biofilm formation (alginate production), effector delivery system (T6SS, T4SS, T2SS), immune modulation (LPS, capsule), motility (flagella, chemotaxis), and nutritional metabolic factors (metal tolerance, siderophores, synthesis of amino acids, nitrogenated bases, and cofactors). For Exoenzymes (mainly enolases and metalloproteases), the highest abundance corresponds to Gemmatimonadota, Actinomycetota, and Pseudomonadota.

Genes encoding putative exotoxins were found in Planctomycetota (Planctomycetales), Bacteroidota, Pseudomonadota, and Actinomycetota. Among these, we found the phytotoxin phaseolotoxin, hemolysins, and cytolysins. For Stress Survival, the highest abundance is observed in Myxococcota (0.5), followed by Actinomycetota (0.44), and Pseudomonadota (0.43), with genes including molecular chaperones, catalases peroxidases, superoxide dismutases, and peroxiredoxins.

Upon expanding our search of ARGs and VFGs to the contigs not included in SRGs, we found additional putative ARGs and VFGs. Besides the gene families already described, we found additional families of beta-lactamases (including TEM, L1, FEZ, VEB, THIN, AST, and OXA), additional aminoglycoside inactivation enzymes including several phosphotransferases, streptogramin acetyltransferases, and additional efflux pumps (Supplementary Table 8).

Among the assembled contigs, we also found additional virulence factors, including the MtrCDE efflux pump, shown to contribute to the survival of *Neisseria gonorrhoeae* from human neutrophils (Handing et al., 2018) (Supplementary Table 9). Also, in terms of exoenzymes, we identified putative orthologs of the exoenzyme hemagglutinin proteinase HapA, a well-characterized factor contributing to *Vibrio cholerae* virulence (Benitez and Silva, 2016), and the alkaline protease AprA, allowing *P. aeruginosa* to block complement activation (Laarman et al., 2012). Additionally, the exotoxins colibactin, highly associated with virulent *K. pneumoniae* and *E. coli* (Faïs et al., 2018; Lam et al., 2018), putative orthologs of the Cya hemolysin adenylate cyclase from *Bordetella* and Haemolysin III from *Bacillus*.

Regarding immune modulation factors, we found putative orthologs of several *Mycobacterium* virulence determinants, including MymA, linked to survival inside activated macrophages and resistance to acidic pH and other harsh conditions (Singh et al., 2022), the Rv1523 rhamnosyl_O-methyltransferase promoting drug resistance and virulence (Ali et al., 2021), the MptpA tyrosine phosphatase responsible for the inhibition of phagosome-lysosome fusion and essential for virulence in this pathogen (Kovermann et al., 2023), and the alkyl hydroperoxide reductase AhpC, involved in counteracting the host’s ROS burst by decomposing hydrogen peroxide into water and oxygen (Wong et al., 2017). Additionally, we found putative orthologs of the MntABC permease influencing *Neisseria gonorrhoeae* growth and interaction with cervical epithelial cells (Lim et al., 2008) and a set of ureases.

### 3.5 Potential association of virulence and resistance factors with mobile genetic elements (MGEs)

We aimed to identify possible virulence or resistance genes associated with MGEs and thus potentially transferrable to other bacteria. For this, we predicted insertion sequences, integrons, and prophages among the assembled contigs and SRGs using dedicated bioinformatic tools. Then, we identified EGMs at a distance of 5 kbp or less from ARGs or VFGs. Among the contigs of the two sites (RU and RD), we identified 20 ARGs and 124 VFGs close to insertion sequences, encompassing 21 families of these mobile elements (Supplementary Figure 3A and Supplementary Table 10). IS5 (17, 10.06%), IS21 (16, 9.47%), and IS110 were the most frequent, along with the ARGs *arr-1*, AAC(3)-Ic, *bacA*, and *bla*_THIN-B_, among others. Also, 81 gene families encoding predicted virulence factors were identified, with putative orthologs of the GDP-mannose 4,6-dehydratase Gmd (*Brucella*), the Tir domain-containing protein BtpB (*Brucella*), and the daunorubicin ABC transporter ATP-binding protein DrrA (*Mycobacterium*) among the most frequent.

We identified three cases of VFGs potentially associated with integron-like elements commonly known as CALINs (clusters of attC sites lacking integron-integrases). One CALIN carried a putative ortholog of the *rfbA/rmlA* gene encoding a glucose-1-phosphate thymidyltransferase involved in rhamnose synthesis and virulence in *Mycobacterium* (Hirmondó et al., 2022) (Supplementary Figure 3B). Of note, this element also carried genes that would originate from distant bacterial lineages, including a monooxygenase from *Halopseudomonas* and a MAPEG family protein from Deltaproteobacteria, supporting the xenogenetic nature of this element. Additionally, we found a putative prophage carrying a gene similar to *csrA*, a key regulator of the virulence and bi-phasic life cycle of *Legionella pneumophila* (Sahr et al., 2017). Sixty-three ARG- or VFG-associated EGMs were located in contigs belonging to SRGs. A higher prevalence of these elements was observed in Actinomycetota SRGs (22), followed by Bacteroidota (16), Pseudomonadota (8), Acidobacteriota (7), Gemmatimonadota (3), Planctomycetota (3), Chloroflexota (2) and Myxococcota (2) (Figure 4).

Since plasmid contigs are commonly lost during the binning process due to their different sequence composition compared to the chromosomal contigs, we conducted a plasmid-directed assembly using metaplasmidSPAdes to identify possible plasmids. 167 and 122 plasmids were assembled from RU and RD metagenomic reads, respectively, with lengths ranging from ∼1.1 to ∼142 kbp. Of them, a predicted ∼16-kbp Actinomycetes plasmid carried putative orthologs of the *glc* genes involved in glyoxylate shunt, a critical process for *Mycobacterium* infection (Huang et al., 2016), along with genes involved in the glycerate metabolism (Supplementary Figure 3C).

Overall, this evidence indicates that, although with a limited frequency, some putative ARGs and VFGs would be associated with mobile elements, pointing out the possible horizontal transference of these genes in these remote and extreme soils.

### 3.6. Bacterial isolates from Union Glacier soil samples

Using four culture media, we obtained 51 bacterial isolates from the Union Glacier soil samples (Supplementary Table 11). Most of them grew in the R2A culture medium (31), followed by LB medium (11), Czapek (5), and NFM (2). Thirty-three isolates came from the RU soil samples, twelve from RD, and four from EH. The isolate classification through full-length 16S rRNA sequencing and analysis revealed four phyla and ten different genera. Actinomycetota accounted for 41 isolates (80%), including six genera from three families of the order Micrococcales, namely, *Arthrobacter* (14), *Mycetocola* (14), *Pseudarthrobacter* (4), *Kocuria* (3), and *Plantibacter* (3). Eight Pseudomonadota isolates were identified: six *Pseudomonas* and two *Sphingomonas*. Also, one *Planococcus* (Bacillota) isolate and one *Flavobacterium* (Bacteroidota).

### 3.7. Antibiotic resistance and virulence factors production among Union Glacier soil bacterial isolates

Using disk diffusion assays adapted to Antarctic soil bacteria (Marcoleta et al., 2022), we tested the sensitivity of the isolates to 35 different clinical antibiotics, covering diverse action mechanisms and chemical nature (Supplementary Table 2). Most *Pseudomonas* isolates stood out for their notable multiresistance, one resisting up to 24 antibiotics (*Pseudomonas* sp. EH124), five to 23, and one to 20 (Figure 5). Other isolates showing multidrug resistance were *Arthrobacter sp.* RD7 (21) and RD8 (20), *Plantibacter sp.* RU18 (20), and *Flavobacterium sp.* EH125 (19). In contrast, other isolates were sensitive to all the antibiotics tested, including some *Arthrobacter*, *Brevibacterium*, *Kocuria, Planococcus, and Plantibacter*. Regarding the antibiotics tested, we observed a high frequency of resistance to beta-lactams, especially aztreonam, ceftazidime, oxacillin, and cefepime. Conversely, no resistant isolates were observed for the cyclines tetracycline, minocycline, and tigecycline. Also, piperacillin, levofloxacin, and ciprofloxacin showed low resistance levels.

**Figure 5.**
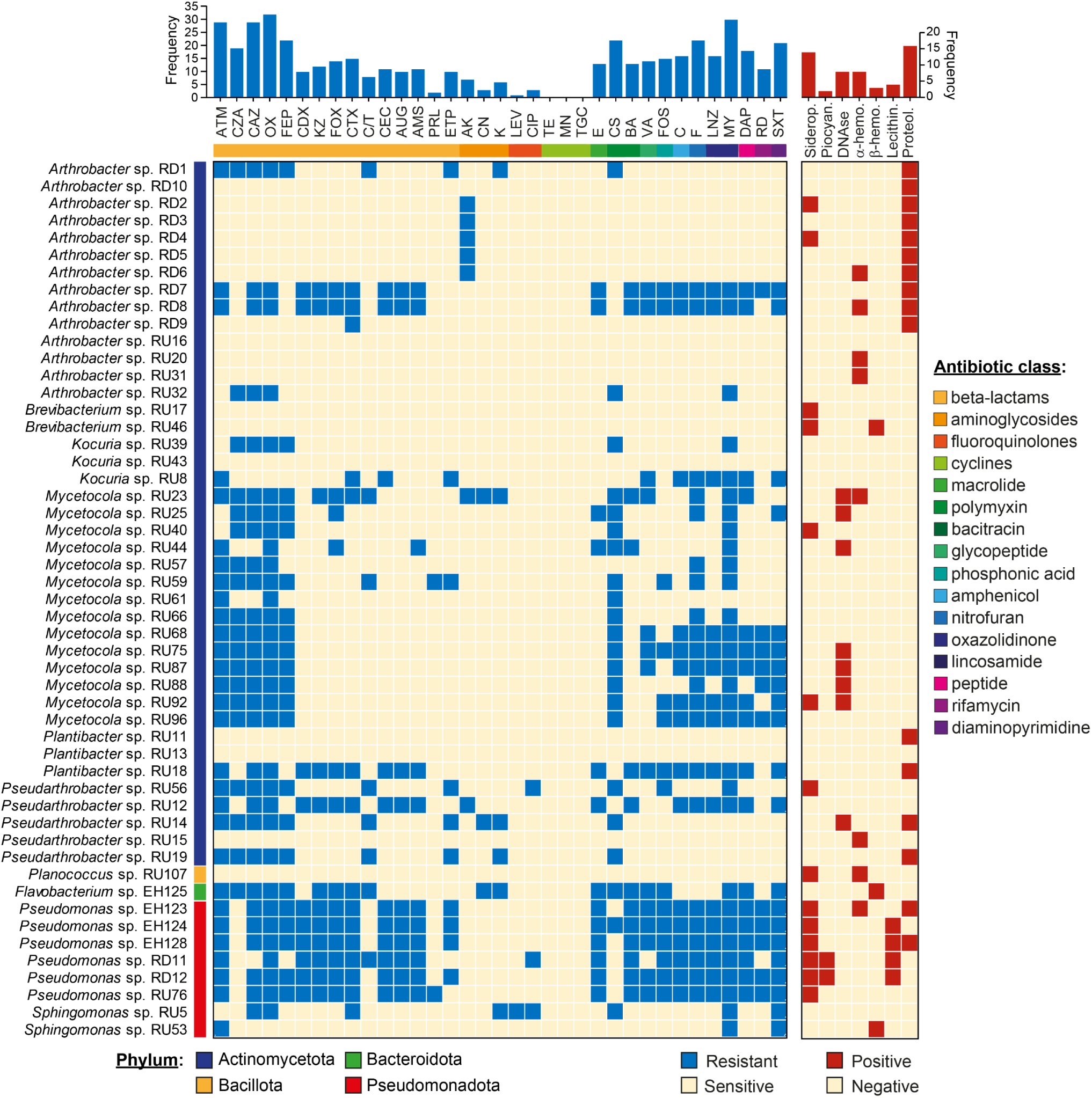
Antibiotic resistance and virulence factor production in Union Glacier bacterial isolates. The antibiotic resistance profile was obtained from disk diffusion assays, while the virulence factor production was evaluated using different indicator agar media, specific for siderophore production and DNAse, hemolytic, lecithinase, and protease activity.

To explore the expression of virulence-related traits among the Union Glacier soil isolates, we adapted chromogenic agar media traditionally used to detect virulence factor production in pathogenic bacteria. Specifically, we used yolk egg agar to detect lecithinase production, an exotoxin targeting the lecithin phospholipids in eukaryotic cells linked to the virulence of some Gram-positive and Gram-negative pathogens. This medium was also used to assess proteolytic activity. Additionally, we used *Pseudomonas* agar to detect the production of pyocyanins, toxic compounds that promote the generation of ROS and subsequent cellular and tissue damage. Furthermore, we used chrome azurol S agar to detect siderophore production, molecules that aid pathogens in scavenging iron during infection. In addition, we used sheep blood agar to detect hemolytic activity.

Most isolates produced one or up to three virulence factors (Figure 5). *Pseudomonas sp.* isolates registered the highest proportion, where they all produced siderophores, most lecithinase activity, and some pyocianins, hemolytic activity, or proteolytic activity. Conversely, *Kocuria* isolates showed no virulence factors. Additionally, several *Mycetocola sp.* isolates showed DNAse activity, while *Arthrobacter* and *Pseudarthrobacter* showed mainly proteolytic activity, and some also showed siderophore production. Also, *Sphingomonas*, *Flavobacterium*, and *Brevibacterium* isolates showed beta-hemolysis. This evidence indicates that part of the microbiota inhabiting Union Glacier soils can produce factors and activities linked to virulence in pathogenic bacteria. Additional studies are required to test the pathogenicity of these bacteria over a host organism.

## 4. DISCUSSION

The melting of Antarctica’s icy landscapes due to global warming would cause previously isolated microbial communities and their genetic content to be exposed, with implications for the ecology and evolution of microbes and global health (Pearce, 2008; Sajjad et al., 2020). Predictions suggest that ice-free areas will triple by the end of the century, highlighting the urgency to understand the impact of climate change on these ecosystems (Lee et al., 2017). Regarding global health, the melting of the Antarctic permafrost could affect the distribution of microbial pathogens and vectors that transmit infectious diseases, exposing humans, animals, and plants to emerging diseases. Here, we showed that even in the hyper-extreme environmental conditions in the Union Glacier cold desert, microbial communities of bacteria, fungi, and archaea are thriving. Moreover, despite this environment isolation and minimal human intervention, some resident bacteria host a relatively ample resistome and virulome, with culturable bacteria displaying multidrug resistance or having pathogenic potential.

The presence of diverse microbial communities in the Union Glacier underscores the resilience and adaptability of microorganisms to harsh conditions. Still, only specific lineages can thrive in this environment, as evidenced by the markedly lower alpha diversity observed (Shannon ranging from ∼3 to ∼5) compared to humanized and non-intervened soils from the Antarctic Peninsula (Shannon ranging from ∼8 to ∼10) (Marcoleta et al., 2022). At the phylum level, the community composition of RU and RD soils was similar to that of other oligotrophic desert communities, with a high abundance of Actinomycetota, Acidobacteriota, with contributions of Pseudomonadota and Bacteroidota (Fierer et al., 2009; Vásquez-Dean et al., 2020). However, the composition of EH soils was strikingly different, with the dominance of Pseudomonadota (mainly *Pseudomonas* spp. and *Acinetobacter* spp.) and Bacillota (mainly *Staphylococcus* spp. and *Bacillus* spp.). This contrasting pattern could be partly explained by the reported ability of members from these genera to resist increased salinity, as detected in EH soils (Egamberdieva et al., 2019; Jeong et al., 2017; Patel et al., 2022; Sand et al., 2011; Schimel et al., 2007; Van Horn et al., 2014). Overall, the dominance of the mentioned phyla agrees with reports from other Antarctic cold desert landscapes, such as the McMurdo Dry Valleys, highlighting their role in supporting fundamental biogeochemical cycles in oligotrophic and extreme environments (Cary et al., 2010; Goordial et al., 2016).

A previous study provided metagenomic insights into the microbiota from Union Glacier soil, focusing mainly on their functional capabilities for carbon, nitrogen, and sulfur cycling (Li et al., 2019). Consistent with our findings, the authors described communities dominated by bacteria, with increased diversity in the Rossman Cove area compared to other studied zones. Pseudomonadota (mainly Gammaproteobacteria) predominated in most samples, although Actinomycetota was also highly abundant, which reinforces our results showing the strong presence of these two phyla in Union Glacier soils. The functional analysis carried out by Li et al. revealed a suite of adaptations to high irradiance and low temperatures. In this context, the presence of Pseudomonadota and Actinomycetota in our samples hints at a community capable of surviving under UV stress and cold conditions, given the well-documented UV resistance and psychrotolerance of many members of these phyla (Jun et al., 2016). These microorganisms’ metabolic versatility and stress resistance mechanisms have implications for biotechnology by offering enzymes and biochemical pathways capable of functioning under extreme conditions.

The fungal diversity in Union Glacier is much more limited and, in agreement with previous reports from Antarctica (Jiya et al., 2023; Sannino et al., 2020; Stoppiello et al., 2023), fifteen out of sixteen ASVs detected in this work belonged to phylum Ascomycota. Most ASVs were affiliated with the Lecanoromycetes class of lichenized fungi and occurred in the three RU samples, which agrees with a previous report of lichens in the Ellsworth Mountains (Øvstedal and Schaefer, 2013), close to Union Glacier. Moreover, due to the characteristic mechanisms that protect lichens from radiation and allow them to withstand freezing temperatures and periods of drought, it is not surprising that Lecanoromycetes have been reported as dominant in cryptoendolithic communities colonizing sandstone rocks in Victoria Land and several other sites in Antarctica (Coleine et al., 2018; Ruprecht et al., 2012; Zucconi et al., 2016). Moreover, members of the *Lecidella*, *Buellia*, and *Acarospora* genera detected in RU samples have been recovered from different Antarctic soil samples (Coleine et al., 2018). In EH soils, we recovered only three ASVs affiliated with Dothideomycetes, a diverse class in the Ascomycota phylum that includes numerous rock-inhabiting fungi (RIF), which can tolerate harsh conditions prevailing on rock surfaces (Coleine et al., 2021, 2018). Besides Ascomycota, a previous work isolated basidiomycetous yeasts from Union Glacier sedimentary rocks (Barahona et al., 2016), although our metabarcoding approach did not detect these taxa in the sampled soils.

The limited but significant detection of archaeal ASVs, particularly from genera *Nitrososphaeraceae* and *Nitrosocosmicus*, indicates the presence of ammonia oxidizers that contribute to the nitrogen cycle in these cold deserts. This finding is consistent with previous reports in polar regions (Alves et al., 2019; Hayashi et al., 2020; Zhang et al., 2020), highlighting the ecological significance of archaea in global nitrogen cycling, even under extreme conditions. Species from *Nitrosocosmicus* were previously reported as prominent ammonia oxidizers in the Arctic (Alves et al., 2019) and in East Antarctica (Hayashi et al., 2020). This study corresponds to the first report of Nitrososphaeraceae and Nitrosocosmicus species in Union Glacier and West Antarctica, suggesting their capacity to thrive in cold and nutrient-deficient conditions.

Although previous studies have explored the microbial diversity of Union Glacier, none of them provided insights into these microbes’s resistome, virulome, and MGEs. In this study, we used shotgun metagenomics to investigate these features and reconstruct species-level representative genomes that will also serve further studies on these microbes’ genomic and functional properties. Previous studies have revealed antibiotic resistance and virulence factors in remote and ancient sites with minimal anthropogenic intervention, showing that they would have evolved along different microorganisms’ competition, predation, antagonism, and survival processes (Allen et al., 2009; D’Costa et al., 2011; Kim et al., 2022; Pawlowski et al., 2016; Wright and Poinar, 2012). Our findings in Union Glacier would correspond to the most meridional report of ARGs and VFGs on our planet, underscoring the ubiquity of antimicrobial resistance and virulence traits in environmental reservoirs. This evidence aligns with the One-Health perspective by demonstrating how environmental microbes, not directly associated with human activities, possess genes that can contribute to the global pool of resistance and virulence factors, potentially impacting public health through horizontal gene transfer mechanisms (Wellington et al., 2013).

Single-strain and metagenomic sequencing combined with phenotypic tests have revealed a remarkable variety of ARGs and antibiotic-resistant bacteria in different areas of the Antarctic Peninsula and the McMurdo Dry Valleys, corresponding to natural resistome from soil, water, and snow, among other sources (Jara et al., 2020; Marcoleta et al., 2022; Ren et al., 2024; Santos et al., 2022; Wei et al., 2015). In most of these sites, a predominance of efflux pumps followed by antibiotic inactivation enzymes and target alteration proteins linked to multidrug, beta-lactam, peptide, aminoglycoside, and vancomycin resistance was observed. These genes were mainly linked to bacteria from the phyla Pseudomonadota, Actinomycetota, and Bacteroidota. Remarkably, similar results were observed in other cryosphere environments, including permafrost from the Arctic (Kim et al., 2022; Xie et al., 2024) and mountain glaciers in Central Asia and the Tibet Plateau (Mao et al., 2023; Ren and Gao, 2024). In agreement, our results showed that bacteria thriving in the Union Glacier, mainly Pseudomonadota and Actinomycetota, host an overall similar array of ARGs. Among the most abundant efflux ARGs were *novA* and *bcrA*, encoding type III ABC transporters of novobiocin (aminocoumarin) and bacitracin (peptide), respectively, and Mex family multidrug pump components. Also, our results and the cited works coincided in a high abundance of the target alteration gene *bacA* linked to bacitracin resistance and in the presence of various genes encoding putative beta-lactamases and aminoglycoside inactivation enzymes. Similarly, vancomycin resistance genes have been found in undisturbed Arctic wetlands (Diaz et al., 2017), soil in the Tibet Plateau (Li et al., 2020), ancient permafrost sediment (D’Costa et al., 2011), and in this work were found in Union Glacier SRGs, mainly from Actinomycetota.

No previous reports have investigated the virulome of Antarctic microbial communities. ARGs and VFGs were investigated in Arctic permafrost in Alaska and in Tibetan glaciers, where adherence, stress tolerance, secretion systems, cell surface, and invasion-related factors were predominant. Further, most of the MAGs recovered corresponding to potentially pathogenic antibiotic-resistant bacteria (PARB) belonged to Actinomycetota and Pseudomonadota, as well as the ARGs and VFGs (Kim et al., 2022; Mao et al., 2023). Concordantly, in the Union Glacier soil metagenomes and SRGs, we observed mainly VFGs linked to secretion systems, stress alleviation, adherence, biofilm formation, capsule, and LPS. Also, hemolysin and cytolysin exotoxins. Whether these environmental virulomes could act as a source of virulence deserves further investigation. In this regard, well-documented cases of VFGs emerging from the environment correspond to some exotoxins from major human pathogens, including the diphtheria toxin of *Corynebacterium diphtheriae*, the Shiga-like toxins of *Escherichia coli*, and the cholera toxin of *Vibrio cholerae*. All of them are encoded in prophages present in different environmental bacteria, which were acquired horizontally by pathogens (Casas et al., 2011; Casas and Maloy, 2018; Friedman and Court, 2001).

The spread of ARGs and VFGs in the environment is mainly attributed to horizontal gene transfer of MGEs encoding them (Bennett, 2008; Partridge et al., 2018), which have been widely recognized as agents promoting bacterial evolution and innovation (Gogarten and Townsend, 2005). Previous evidence indicates that the ARGs actively transferred between bacterial populations by HGT correspond mainly to those coding for enzymes that allow antibiotic inactivation (Nielsen et al., 2022) and antimicrobial efflux pumps (Sánchez-Osuna et al., 2023), which are of great clinical relevance. Furthermore, a recent global survey of metagenomes from different habitats indicated that roughly 24% of the ARGs found would pose a health risk for humans (Zhang et al., 2022). Thus, it is relevant to explore the association of environmental ARGs and VFGs with MGEs, and the potential for transferring these determinants to human-associated bacteria. In Union Glacier soil, we detected limited cases of ARGs or VFGs associated with mobile elements, which agrees with the restricted microbial diversity found and a potentially low lateral gene flow. Nevertheless, examples of insertion sequences, integron-like elements, and prophages close to these genes were identified. Evidence from McMurdo (Wei et al., 2015), Antarctic Peninsula (Santos et al., 2022), Arctic permafrost (Kim et al., 2022), and other glacier environments (Wang et al., 2024) indicate that prophages would be the main actors in ARG and VFG horizontal transference. Although our approach identified a limited number of prophages in Union Glacier, further studies targeting the virome of this area would provide a clearer view of its implications in this ecosystem.

Although investigating the resistome and virulome in the environment has benefited significantly from the development of metagenomic approaches, as an essential drawback, these methodologies allow predictive phenotypic determinations that must be validated experimentally. In this direction, we isolated several Union Glacier bacterial strains from the genus *Pseudomonas*, *Mycetocola*, *Pseudarthrobacter*, and *Arthrobacter*, displaying resistance to different antibiotic classes, some also producing virulence factors, which agrees with our metagenomic findings. Previous reports described autochthonous multiresistant Antarctic bacteria, including *Pseudomonas*, *Arthrobacter*, *Pedobacter*, and Acinetobacter (Marcoleta et al., 2022; Opazo-Capurro et al., 2019; Tam et al., 2015). Interestingly, within our isolates, we encountered multiresistant *Arthrobacter* and *Pseudomonas* isolates producing virulence factors such as proteases and siderophores. These two genera would be significant hosts for ARGs and VFGs in Antarctica. Further investigation is warranted to determine if the production of these virulence factors confers an adaptive advantage for survival in the challenging conditions of the Antarctic desert. Moreover, additional studies using infection models are required to evaluate the virulence of these isolates over different hosts.

In sum, we unveiled the diversity, structure, resistome, and virulome of microbial communities thriving in the hyper-extreme Union Glacier cold desert, evidencing that, although with limited abundance, a variety of putative ARGs and VFGs are present, some associated with mobile elements. These results and others from remote and extreme ecosystems emphasize the need for integrated environmental and public health surveillance to address AMR. Although the low frequency of ARGs and VFGs associated with MGEs would limit the Union Glacier resistome and mobilome as a source of genes transferred to clinical pathogens, this environment hosts potentially pathogenic antibiotic-resistant bacteria, as evidenced by some of the isolates. Further studies are required to address their relative risk.

## 5. Data availability

The 16S rRNA sequences of bacterial isolates were submitted to GenBank. Accession numbers are provided in Supplementary Table 3.

The metabarcoding reads were submitted to NCBI under the BioProject PRJNA1067332.

The shotgun Illumina and Nanopore metagenomic reads were submitted to the NCBI Sequence Read Archive (SRA) under the accessions SRR28324393, SRR28324394, and SRR28324392 (BioProject PRJNA1067332).

The species-level representative genomes were submitted to the NCBI Genome database under the BioProject PRJNA1067332. Individual genome accession numbers are provided in Supplementary Table 5.

## 6. Acknowledgments

This study was funded by grants mBioClim ACT210044, Polarix ACT192057, and FONDECYT 1221143 from Agencia Nacional de Investigación y Desarrollo (ANID), Chile.

