## Supplementary Figures and Tables for "Life on the edge: microbial diversity, resistome, and virulome in soils from the Union Glacier cold desert": Arros_et_al_Supplementary_Figures_and_Tables.pdf

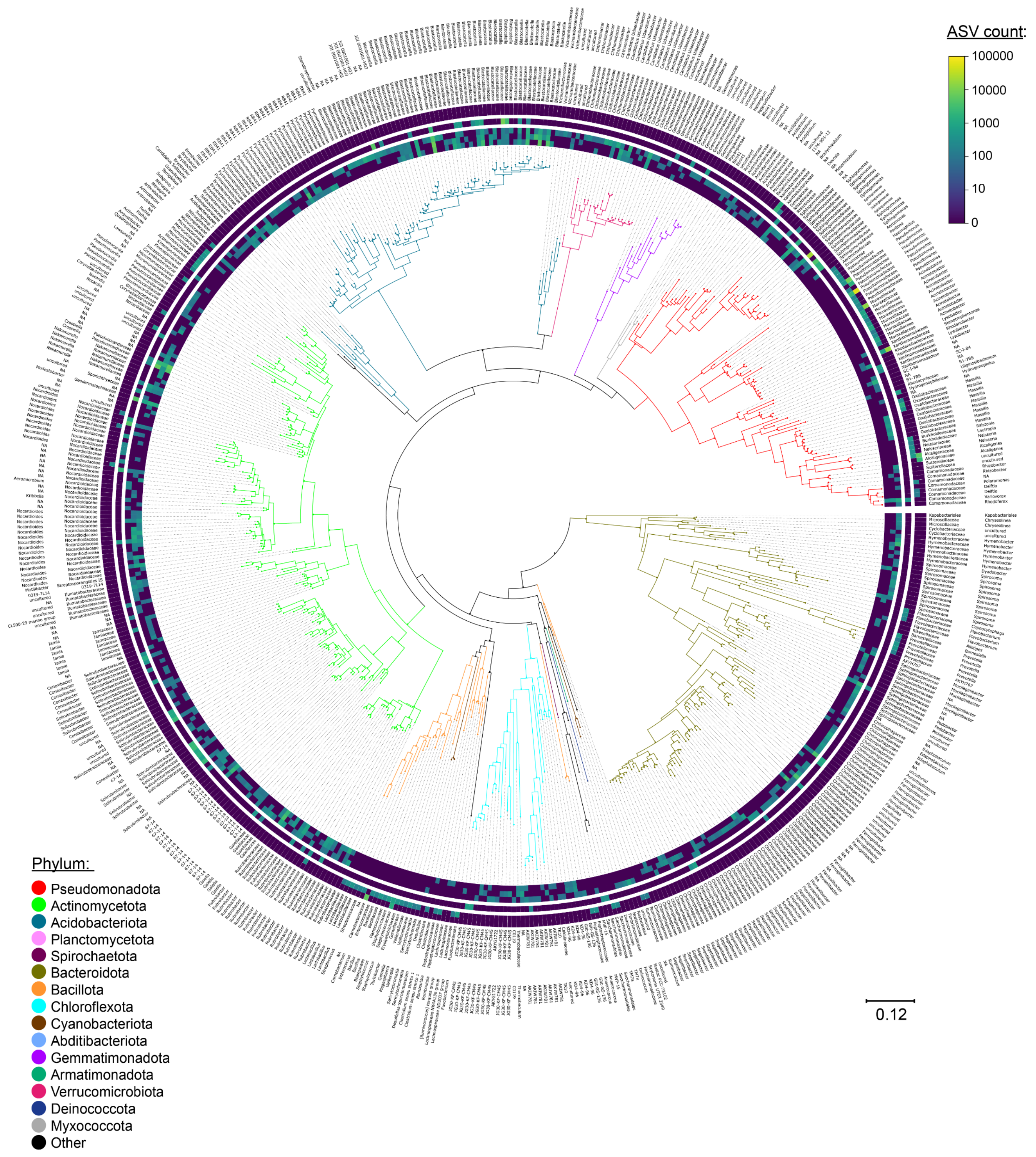


**Supplementary Figure 1. Bacterial ASVs detected in Union Glacier soils.** Maximum-likelihood tree showing the phylogenetic relationships among the bacterial ASVs. The colors in the branches represent the different phyla. From inner to outer, the tracks indicate the relative abundance of each ASV in RU1, RU2, RU3, RD, EH1, and EH2 soil samples. Also, the family and the genus are indicated for each ASV.

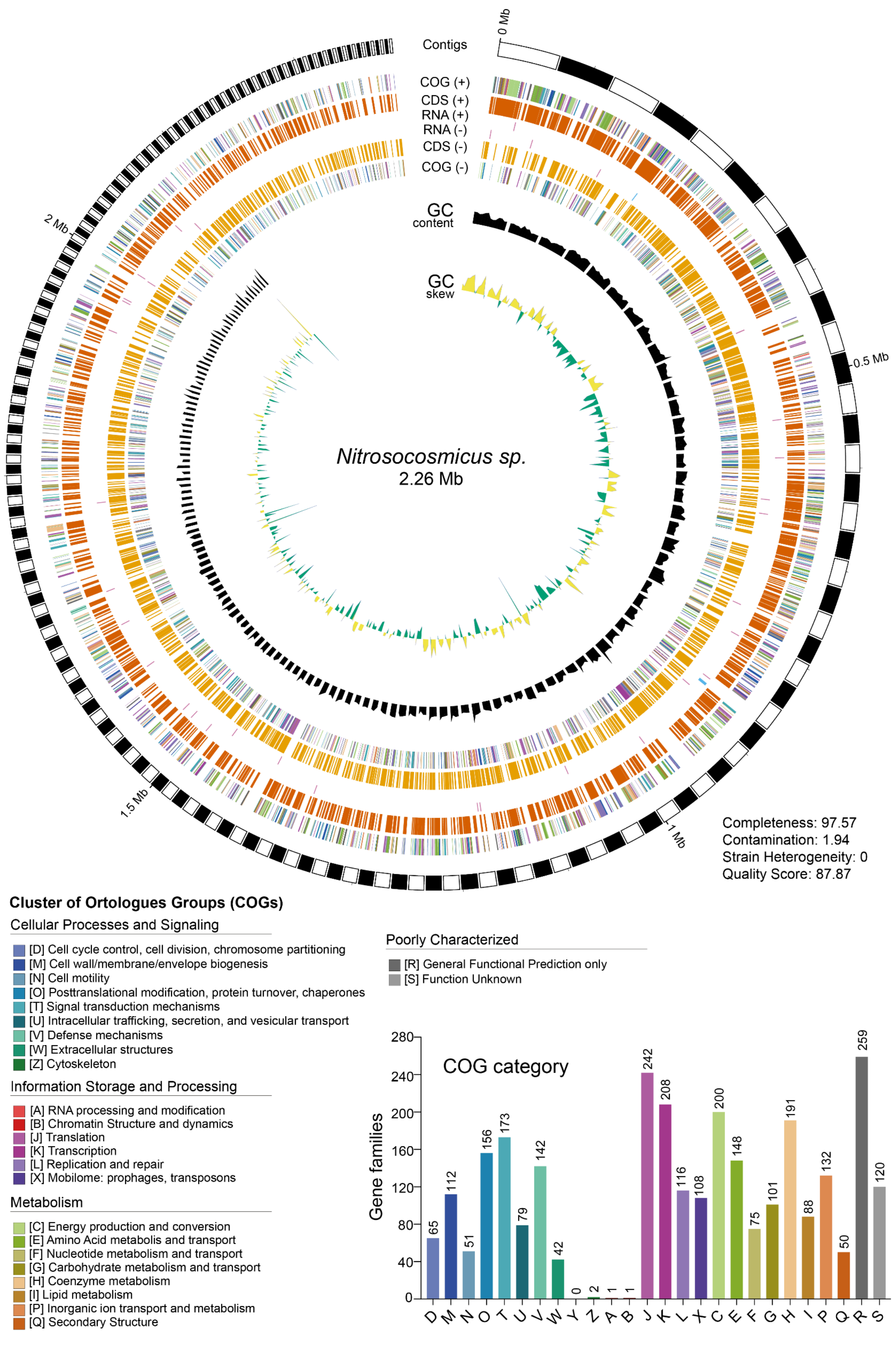


**Supplementary Figure 2. Species-level representative genome of the archaeon *Nitrosocosmicus sp.* recovered from Union Glacier soil.** The circular map tracks depicts, from inner to outer, the GC skew and content, the COGs, CDSs, and RNAs encoded in the negative strand, followed by the RNAs, CDSs and COGs encoded in the positive strand. Relevant genome parameters are also provided.

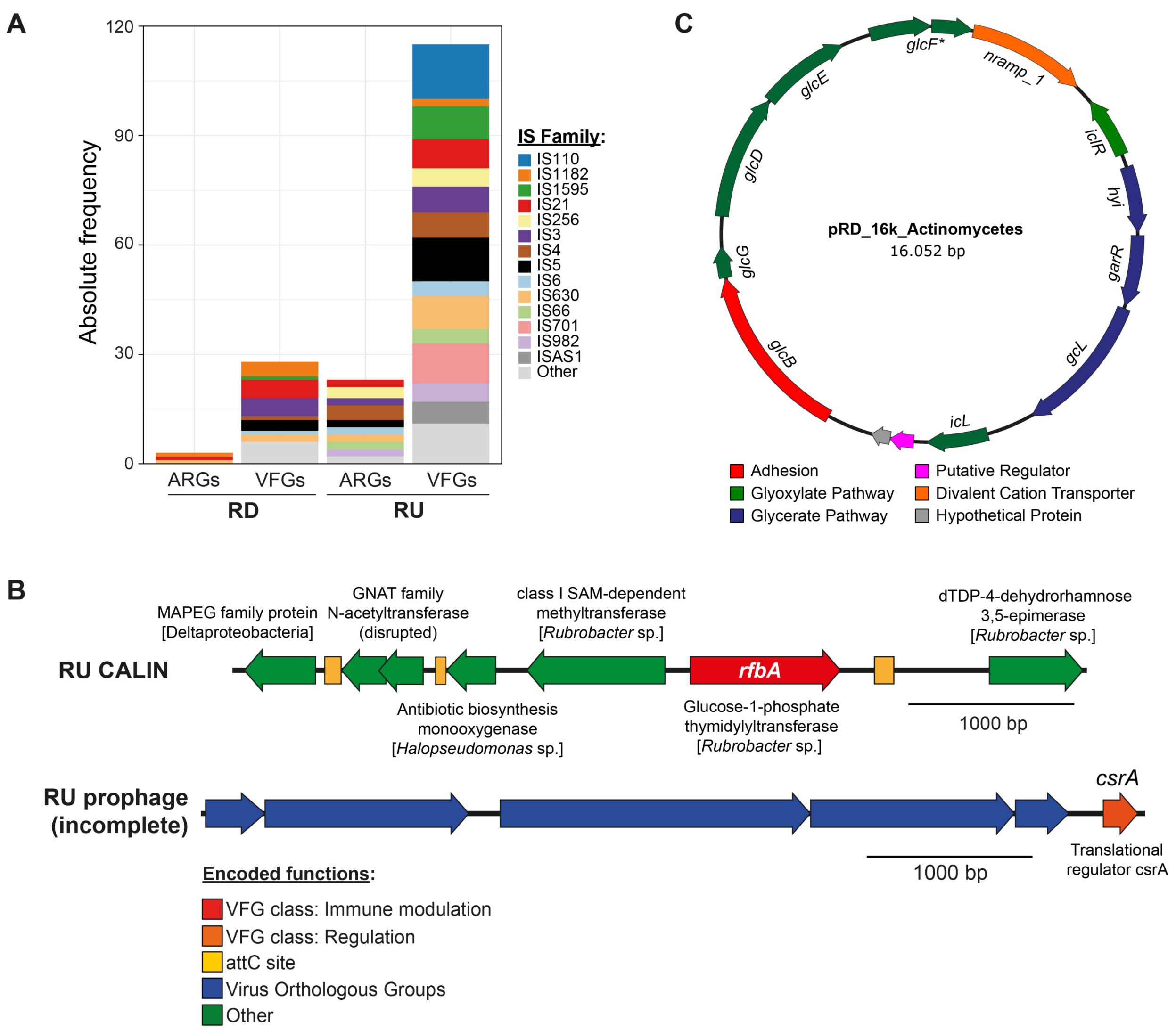


**Supplementary Figure 3. Putative association between mobile genetic elements and VFGs or ARGs.** (A) Different families of insertion sequences located at a distance of 5 kbp or less from one or more VFGs and ARGs. Only IS families with an absolute frequency of six or more are individualized (B) putative CALIN and prophage associated with the *rfbA* and *csrA* genes, respectively, found in contigs from the RU soil metagenome. (C) Predicted plasmid from Actinomycetes carrying the *glcB* gene linked to adhesion in *Mycobacterium*, assembled from the RD soil metagenome.

**Supplementary Table 1. Physicochemical properties of soil samples from Union Glacier**

| Sample | TP ($mg\ kg^{-1}$) | $P_{Olsen}$ ($mg\ kg^{-1}$) | TN ($mg\ kg^{-1}$) | TC ($mg\ kg^{-1}$) | pH | EC ($\mu s\ cm^{-1}$) | OM (%) |
|---|---|---|---|---|---|---|---|
| **RD** | 101.27 | 7.16 | 1.83 | 3.00 | 8.4 | 355 | 0.29 |
| **RU1** | 96.63 | 7.19 | 2.21 | 6.64 | 8.2 | 171 | 0.38 |
| **RU2** | 124.41 | 11.06 | 3.39 | 10.38 | 8.0 | 89 | 0.72 |
| **RU3** | 200.68 | 8.80 | 1.89 | 4.55 | 8.2 | 39 | 0.32 |
| **EH1** | 119.94 | 2.72 | 2.07 | 11.73 | 7.7 | 1535 | 1.00 |
| **EH2** | 133.35 | 6.70 | 3.36 | 17.82 | 7.8 | 2067 | 1.30 |

The analyzed samples were the base of Mount Rossman (RD), Rossman Cove summit (RU1-RU3), and Edson Hills summit (EH1-EH2).

**Supplementary Table 2.** Antibiotics used for disk diffusion assays.

| Abbreviation | Antibiotic | Antibiotic class | Antibiotic subclass | Antibiotic mass in the disk (ug) |
|---|---|---|---|---|
| AK | Amikacin | Aminoglycosides | Aminoglycosides | 30 |
| CN | Gentamicin | Aminoglycosides | Aminoglycosides | 10 |
| K | Kanamycin | Aminoglycosides | Aminoglycosides | 30 |
| C | Chloramphenicol | Amphenicols | Amphenicols | 30 |
| ETP | Ertapenem | Beta-lactams | Carbapenems | 10 |
| CDX | Cefadroxil | Beta-lactams | Fifth-generation cephalosporins | 30 |
| KZ | Cefazolin | Beta-lactams | Fifth-generation cephalosporins | 30 |
| C/T | Ceftolozane-tazobactam | Beta-lactams | Fifth-generation cephalosporins | 40 |
| FEP | Cefepime | Beta-lactams | Fourth-generation cephalosporins | 30 |
| ATM | Aztreonam | Beta-lactams | Monobactams | 30 |
| OX | Oxacillin | Beta-lactams | Penicillins | 1 |
| PRL | Piperacillin | Beta-lactams | Penicillins | 100 |
| CEC | Cefaclor | Beta-lactams | Second-generation cephalosporins | 30 |
| FOX | Cefoxitin | Beta-lactams | Second-generation cephalosporins | 30 |
| CTX | Cefotaxime | Beta-lactams | Third-generation cephalosporins | 30 |
| CAZ | Ceftazidime | Beta-lactams | Third-generation cephalosporins | 30 |
| CZA | Ceftazidime-avibactam | Beta-lactams | Third-generation cephalosporins | 50 |
| AUG | Amoxicillin-clavulanic acid | Beta-lactams plus beta-lactamase-inhibitor | Penicillins plus beta-lactamase inhibitor | 30 |
| AMS | Ampicillin-sulbactam | Beta-lactams plus beta-lactamase-inhibitor | Penicillins plus beta-lactamase inhibitor | 30 |
| CIP | Ciprofloxacin | Fluoroquinolones | Fluoroquinolones | 5 |
| LEV | levofloxacin | Fluoroquinolones | Fluoroquinolones | 5 |
| VA | Vancomycin | Glycopeptides | Glycopeptides | 30 |
| TGC | Tigecycline | Glycylcyclines | Glycylcyclines | 15 |
| MY | Lincomycin | Lincosamides | Lincosamides | 2 |
| DAP | Daptomycin | Lipopeptides | Lipopeptides | 30 |
| E | Erythromycin | Macrolides | Macrolides | 15 |
| F | Nitrofurantoin | Nitrofuran-derivatives | Nitrofuran-derivatives | 300 |
| LNZ | Linezolid | Oxazolidinones | Oxazolidinones | 30 |
| BA | Bacitracin | Peptide | Polypeptide | 10 |
| CS | Colistin sulfate | Peptide | Polymyxins | 10 |
| FOS | Fosfomycin | Phosphonics | Phosphonics | 200 |
| RD | Rifampicin | Rifamycins | Rifamycins | 5 |
| SXT | Trimethoprim-sulfamethoxazole | Sulfonamide-trimethoprim | Sulfonamide-trimethoprim | 25 |
| MN | Minocycline | Tetracyclines | Tetracyclines | 30 |
| TE | Tetracycline | Tetracyclines | Tetracyclines | 30 |

**Supplementary Table 3.** Bacterial, fungal, and archaeal ASVs observed in Union Glacier soil samples (provided as a separate spreadsheet).

**Supplementary Table 4.** Union Glacier soil metagenome sequencing and assembly stats.

| | Sample/assembly | | | |
|---|---|---|---|---|
| | RDP | RUP | | |
| | Illumina only | Illumina only | ONT only | Hybrid |
| Nanopore total bases (Gbp) | - | - | 16.4 | 16.4 |
| Nanopore average read length (bp) | - | - | 6995.2 | 6995.2 |
| Nanopore average read quality (Q) | - | - | 12.2 | 12.2 |
| Illumina total bases (Gbp) | 15.8 | 19.0 | - | 19.0 |
| Illumina average read length (bp) | 140 | 140 | - | 140 |
| Illumina average read quality (Q) | 36 | 36 | - | 36 |
| Nonpareil diversity | 19.68 | 21.11 | - | - |
| Estimated coverage (%) | 88.2 | 79.2 | - | - |
| Total assembly length (Mbp) | 404.5 | 477.9 | 1239.4 | 1100.7 |
| # assembled contigs | 158,748 | 214,816 | 39,069 | 289,300 |
| Largest contig (Mbp) | 0.4 | 0.5 | 4.5 | 1.1 |
| GC content (%) | 63.5 | 56.7 | 62.0 | 59.4 |
| N50 (bp) | 2970 | 2316 | 56,797 | 5590 |
| L50 (contigs) | 30,014 | 45,515 | 5200 | 46,608 |

**Supplementary Table 5.** Main features of the species-level representative genomes (SRGs) recovered from Union Glacier soil metagenomes (provided as a separate spreadsheet).

**Supplementary Table 6.** Putative antibiotic resistance genes identified among the species-level representative genomes (SRGs) recovered from Union Glacier soil metagenomes (provided as a separate spreadsheet).

**Supplementary Table 7.** Putative virulence genes identified among the species-level representative genomes (SRGs) recovered from Union Glacier soil metagenomes (provided as a separate spreadsheet).

**Supplementary Table 8.** Putative antibiotic resistance genes identified among the assembled contigs from Union Glacier soil metagenomes (provided as a separate spreadsheet).

**Supplementary Table 9.** Putative virulence genes identified among the assembled contigs from Union Glacier soil metagenomes (provided as a separate spreadsheet).

**Supplementary Table 10.** Predicted mobile genetic elements comprising or in close proximity to putative antibiotic resistance or virulence genes (provided as separate spreadsheet).

**Supplementary Table 11.** Bacterial isolates recovered from Union Glacier soil samples (provided as separate spreadsheet).
